# Lipid Lowering Oxopropanylindole Hydrazone Derivatives with Anti-oxidant and Anti-hyperglycemic Activity

**DOI:** 10.1101/477315

**Authors:** Kanika Varshney, Amit K. Gupta, Ravi Sonkar, Salil Varshney, Akanksha Mishra, Geetika Bhatia, Anil Gaikwad, Arvind Kumar Srivastava, Mridula Saxena, Sudha Jain, Anil K.Saxena

**Author notes:** Address correspondence to this author at the Medicinal & Process Chemistry Division, Central Drug Research Institute, Lucknow, 226031, India, /Fax: **, +*91-52226123405*; E-mails: (AKS); (SJ).

## Abstract

A series of substituted oxopropanylindole hydrazone derivatives was synthesized and evaluated for anti-oxidant and anti-dyslipidemic activity. Among these 12 compounds the three compounds **6c**, **7b** and **7d** showed good anti-oxidant activity and the compound **6c** attenuataed LDL oxidation by 32%. The compounds **6c** and **7d** also showed good anti-dyslipidemic activity by reducing serum levels of total cholesterol (TC), phospholipids (PL) and triglycerides (TG). These two compounds were further evaluated for anti-adipogenic and anti-hyperglycemic activity, where **6c** was found most active compound with 44% reduction in lipid accumulation and 20.5% and 24.3% reduction in blood glucose at 5h and 24h respectively, as compared to standard drug metformin.

## 1. Introduction

Atherosclerosis is a key contributor in the cardiovascular diseases (CVDs) which are the major cause of mortality around the world. According to WHO, by 2030 more than 23 million people will die annually from CVDs [1]. These are multifactorial diseases. Diseases like obesity, diabetes mellitus, impaired glucose tolerance, insulin resistance, dyslipidemia (high LDL cholesterol, low HDL cholesterol, and high triglyceride levels in blood), and increased blood pressure raise the risk of cardiovascular diseases [2].

The involvement of oxidative stress in combination with hyperlipidemia in the pathogenesis and development of atherosclerosis as well as type 2 diabetes has recently been reported [3]. There is growing evidence that hydroxyl free radicals are involved as major factor causing peroxidative damage to lipoproteins present in the blood, which are responsible for the onset and advancement of atherosclerosis [4]. Moreover, in hyperglycemic patients, the occurrence of several non-enzymatic glycosylations is accompanied by glucose oxidation catalyzed by Cu2+ and Fe2+ resulting in the formation of O2- and OH• radicals which further accelerates the risk of cardiac diseases in dyslipidemic patients [5].

Furthermore, the hallmarks of type2 diabetes viz. the chronic hyperglycemia, insulin resistance and abnormal lipoprotein profiles, contribute in decreasing the bioavailability of vascular nitric oxide (NO), an endothelium derived relaxing factor, and thus impairs the endothelium which increases the risk of atherosclerosis [4].

Therefore, a dyslipidemic agent lowering the cholesterol along with anti-oxidant activity and anti-diabetic activity will be able to protect endothelial and myocardial function and thus may serve as a good anti-atherosclerotic agent. The drugs used in the treatment of dyslipidemia, act by lowering cholesterol or by lowering triglyceride levels in plasma [7–10]. The commonly used anti-dyslipidemic statins have several side effects like myostitis, arthralgias, gastrointestinal upset and elevated liver function tests. Therefore, there is a need to discover potentially better anti-dyslipdemic agents.

Hydrazones as well as carboxamides represent an important class of compounds found in versatile building blocks for the preparation of pharmaceuticals [11, 12]. Moreover, these classes of compounds have been clinically used as therapeutic options in the treatment of diabetes, obesity, metabolic syndrome (dyslipidemia) and CVDs [12, 13].

Several tryptophan derivatives are known to function as a free radical scavenger and anti-oxidants by stimulating several antioxidative enzymes and stabilize cell membranes that help to resist free radical damage [14, 15]. Considering this background, it appeared of interest to synthesize a series of oxopropanylindole hydrazone derivatives and evaluate them against anti-oxidant, anti-dyslipidemic and anti-hyperglycemic activities. Synthesis and biological (anti-oxidant, anti-dyslipidemic and anti-hyperglycemic) activities of a series of 12 substituted oxopropanylindole hydrazone derivatives with different functionalities at R_1_ and R_2_ are reported in this paper.

## 2. Chemistry

The synthesis of target compounds is outlined in Scheme 1. The starting material DL-tryptophan methyl ester (**1**) was synthesized by our earlier reported procedure [16] which on reaction with substituted benzoyl chlorides in the presence of triethylamine afforded respective N-benzoyl tryptophan methyl esters (**2**-**4**). These esters on refluxing with hydrazine hydrate in ethanol gave the key intermediates **5**-**7**. Condensation of key intermediates with substituted benzaldehydes gave the desired compounds (**5a**-**5d**, **6a**-**6d**, **7a**-**7d**). (Scheme 1)

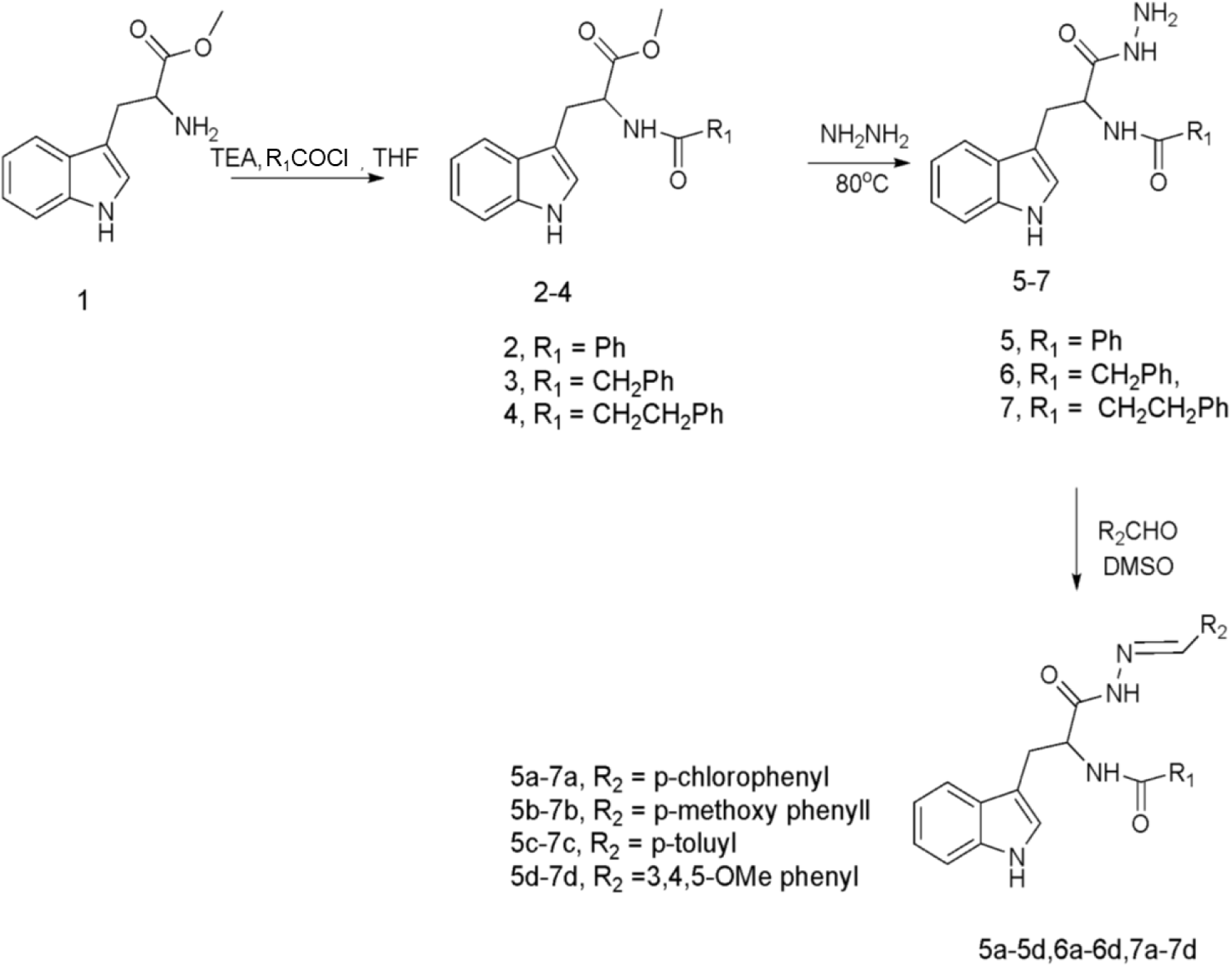

**Scheme 1**

## 3. Materials and Methods

### 3.1. Animal used

Rats were kept in a room with controlled temperature at 25-26 ºC, humidity 60-80%- and 12/12-hours light/dark cycle (light on from 8.00 AM to 8 PM) under hygienic conditions. Animals were acclimatized for one week before starting the experiment. The animals had free access to the normal diet and water. The animal studies and experimental protocols were approved by the institutional ethics committee of CDRI, Lucknow which strictly follows the guidelines of the Committee for Control and Supervision of Experiments on Animals (CPCSEA).

### 3.2 Anti-oxidant activity

Super oxide anions were generated enzymatically by xanthine (160 mM), xanthine oxidase (0.04 U), and nitroblue tetrazolium (320 µM) in absence and presence of the test compounds listed in Scheme 1 at concentrations 200 µg/mL in 100 mM phosphate buffer (pH 8.2). Compounds were sonicated well in phosphate buffer before use. The reaction mixtures were incubated at 37ºC and after 30 min the reaction was stopped by adding 0.5 mL glacial acetic acid. The amount of formazone formed was calculated spectrophotometrically. In another set of experiments, the effect of compounds on the generation of hydroxyl radical was also studied by non-enzymatic reactants. Briefly, •OH were generated in a non-enzymatic system comprising deoxy ribose (2.8 mM), FeSO_4_.7H_2_O (2 mM), sodium ascorbate (2.0 mM) and H_2_O_2_ (2.8 mM) in 50 mM KH_2_PO_4_ buffer (pH 7.4) to a final volume of 2.5 mL. The above reaction mixtures in the absence or presence of test compounds were incubated at 37 °C for 90 min. The test compounds were also studied for their inhibitory action against microsomal lipid peroxidation in vitro by nonenzymatic inducer. Reference tubes and reagents blanks were also run simultaneously. Malondialdehyde (MDA) contents in both experimental and reference tubes were estimated spectrophotometrically by thiobarbituric acid [17]. Alloprinol, mannitol and α-tocopherol were used as standard drugs for superoxide, hydroxylations and microsomal lipid peroxidation.

### 3.3 LDL oxidation

Serum was separated from the blood of normolipidemic donors who were fasted overnight and fractionated into very low-density lipoprotein (VLDL), low density lipoprotein (LDL) and high-density lipoprotein (HDL) by ultracentrifugation [18]. The lipoproteins preparations were dialyzed against 150 mM NaCl containing EDTA (0.02 % w/ v) in presence of N_2_ gas in cold. The purity of LDL was checked on polyacrylamide gel electrophoresis. LDL (0.71 mg) and CuCl_2_.2H_2_O (10 µM) in the absence or presence of the test compounds in 50 mM phosphate buffer saline (pH 7.4) to a final volume of 1.5 ml, was incubated at 37°C for 16 hours. The level of lipid peroxides in unoxidized LDL, oxidized LDL with Cu^++^ in the absence or presence of test compounds at dose of 200 µg/mL were assayed as thiobarbuteric acid reactive substances (TBARS). Briefly the reaction mixture contained 0.5ml SDS (8% w/v), 0.5 ml glacial acetic acid, 1.5 ml TBA (0.8% w/v) was heated in a boiling water bath for one hour. After cooling up to room temperature optical density of reaction mixture was read at 532 nm with respective reagent blank. The level of lipid peroxide as nmol of Malondialdehyde formed was calculated by taking absorption Coefficient of MDA as 1.78×105 cm^-1^ M^-1^ mg protein.

### 3.4 Anti-dyslipidemic activity in triton induced hyperlipidemic rat model

Rats were divided into different groups-control, triton treated and triton plus compounds treated groups, every group containing six rats each. In the acute experiment of 18 hours, hyperlipidemia was developed by administration of triton WR-1339 (Sigma Chemical Company, St. Louis, MO, USA) at a dose of 400 mg/kg, b.w. intraperitoneally (i.p) to animals of all the treated groups with extraction, fractionation and the test compounds were macerated with gum acacia, suspended in water and fed simultaneously with triton at a dose of 100 mg/kg p.o. to the animals of treated groups, diet was withdrawn. Animals of control and triton group without treatment with extraction, fractionation and the test compounds were given same amount of gum acacia suspension (vehicle). After 18 hours of treatment the animals were anaesthetized with thiopentone solution (50 mg/kg b.w.) prepared in normal saline and 1ml blood was withdrawn from retro-orbital sinus using glass capillary in EDTA coated tubes (3.0 mg/ml blood). The blood was centrifuged at 2500 x g for 10 minutes at 4°C and plasma was separated. Plasma was diluted with normal saline (ratio of 1:3) and used for analysis of total cholesterol (TC), Phospholipid (PL) and triglyceride (TG) by standard enzymatic methods. Using Beckmann auto-analyzer and standard kit purchase from Merck Company and post-heparin lipolytic activity (PHLA) was assayed using spectrophotometer [19].

### 3.5 Lipoprotein measurement

Plasma was fractionated into very low-density lipoprotein (VLDL), low density lipoprotein (LDL) and high-density lipoprotein (HDL) by poly anionic precipitation methods. Lipoproteins were analyzed for their triglyceride (TG) level by standard procedures reported earlier [20].

### 3.6 Lipoprotein lipase activity in liver of triton induced hyperlipidemic rats

Liver was homogenized (10%, w/v) in cold 100 mM phosphate buffer pH 7.2 and used for the assay of total lipolytic activity of lipoprotein lipase (LPL) [18].

### 3.6 Molecular modeling

Human lipoprotein lipase (Uniprot Seq: P06858) plausible structural model was built using the single template of horse lipoprotein lipase protein (PDB ID: 1HPL) using Swiss model server. Sequence identity between Human and horse lipoprotein lipase protein is around ~30 % which makes is good candidate for template selection [21]. Homology model was inspected to ensure that the side chains of the conserved residues were aligned to the template. The outlier residues were examined using the Ramachandran plot and the refined using Prime module of Schrodinger software package. Protein and ligands were prepared using default protocol implemented in Schrodinger software package [22]. Compounds were subjected to the InducedFit docking algorithm of the Schrodinger software package where the amino acid residues of proteins were also allowed to move freely along with docked ligand. The receptor grid (LPL binding site) was defined as reported previously [23]. The lowest energy conformation among the largest populated cluster sharing common interactions was selected as the best binding pose for docked ligands.

### 3.7 Cell culture and adipogenic differentiation

3T3-L1 mouse embryo fibroblasts cell line was obtained from the American Type Culture Collection. Cells were cultured in a humidified atmosphere at 37°C and 5% CO2 in Dulbecco’s modified Eagle’s medium (DMEM) containing 10% (v/v) heat-inactivated fetal bovine serum and antibiotic penicillin and streptomycin. For adipogenesis induction 50000 cells were seed in 24 multi-well plates. After 2 days, when cells achieved nearcomplete confluence, culture media was replaced with adipogenesis media I (containing Insulin 5 µg/ml, IBMX 0.5 mM and Dexamethasone 250 nm in culture medium). This media was then replaced after 72 hours with adipogenesis media II (Insulin 5 µg/ml in DMEM with 10% FBS). After replacement of this media, cells were then maintained next 2 days in 10% FBS containing DMEM medium. Lipid globules in the adipogenic cells starts forming from day 4th onwards after treatment, and fully developed adipocytes were observed after day 8th of adipogenesis treatment. More than 80% cells do have lipid globules at this stage.

### 3.8 Triglyceride assay and Oil Red O staining

To study effect of compound on adipogenic differentiation, cells were differentiated as mentioned in above protocol along-with compound at 20 µM concentrations. Fully differentiated 3T3-L1 (with or without compound) adipocytes were rinsed in phosphate buffered saline (pH 7.4). The adipocytes lipid globules were stained with Oil Red O (0.36% in 60% Isopropanol) for 20 min. Unstained Oil Red O was removed by rinsing wells twice with phosphate buffer saline. After complete removal of PBS, finally, 100% Isopropanol was used to extract the dye from the cells and extracted dye absorbance was measured at 492 nm.

### 3.9 Anti-hyperglycemic activity

Male albino rats of Sprague-Dawley strain (8 to 10 weeks of age body weight 160 ± 20 g) were selected for this study. Streptozotocin (Sigma, USA) was dissolved 100 mM citrate buffer pH 4.5 and calculated amount of the fresh solution was injected to overnight fasted rats (60 mg/kg) intraperitoneally. Fasting blood glucose was checked 48 h later by glucometer by using glucostrips and animals showing blood glucose values over 270 mg/dl were selected and divided into groups of five animals each. Rats of experimental groups were administered suspension of standard drug and the desired test samples orally (made in 1.0% gum acacia) at a dose of 100 mg/kg body weight. Animals of control group were given an equal amount of 1.0 % gum acacia. The blood glucose level of each animal was determined just before the administration of standard drug and test samples (0 min) and thereafter at 30, 60, 90, 120, 180, 240, 300 and 1440 min. Food but not water was withdrawn from the cages during 0 to 300 min. The average lowering in blood glucose level between 0 to 300 min and 0 to 1440 min was calculated by plotting the blood glucose level on y-axis and time on x-axis and determining the area under curve (AUC). Comparing the AUC of experimental group with that to control group determined the percent lowering of blood glucose level during the period. Statistical analysis was made by Dunnett’s test (Prism Software).

### 3.10 Statistical evaluation

All results are presented as the means ± S.D. of results from three independent experiments. Groups were analyzed via t-tests (two-sided) or ANOVA for experiments with more than two subgroups. Probability values of p < 0.05 were statistically significant

## 4 Results and Discussion

### 4.1 Anti-oxidant activity

Anti-oxidant activity of test compounds were evaluated by measuring the scavenging potential at 200 µg/ml against formation of O_2_- and OH^2^ in non-enzymatic systems listed in Table 1. Further, the effect of compounds on lipid peroxidation in microsomes were also studied. Out of total 12 compounds screened three compounds **6c**, **7b**, and **7d** showed significant decrease in superoxide anions (33%, 27%, and 25%) as compared to the standard drug allopurinol which showed 43% inhibition of superoxide anions. The compounds **6c**, **7b**, and **7d** also reduced hydroxyl radicals by 35%, 29%, and 23% respectively in comparison to mannitol which showed 47% inhibition in hydroxyl radicals.

**Table 1.** Effect of compounds on generation of superoxide anions, hydroxyl radicals and lipid-peroxidation.

| Series of compounds | Dose (µg/ml) | Superoxide anions (%) (n mol formazone formed/minute) | Hydroxyl radicals (%) (n mol MDA formed/h/mg protein) | Lipid-peroxidation (%) (n mol MDA formed/h/mg protein) |
| --- | --- | --- | --- | --- |
| 5a | 200 | -21** | -20*** | -21** |
| 5b | 200 | -16* | -18* | -15* |
| 5c | 200 | -11* | -10* | -12* |
| 5d | 200 | -20** | -20** | -22** |
| 6a | 200 | -17* | -19* | -19* |
| 6b | 200 | -11* | -13* | -14* |
| 6c | 200 | -33*** | -35*** | -34*** |
| 6d | 200 | -14* | -12* | -13* |
| 7a | 200 | -17* | -15* | -19* |
| 7b | 200 | -27*** | -29*** | -26*** |
| 7c | 200 | -14* | -17* | -15* |
| 7d | 200 | -25*** | -23*** | -23*** |
| Standards | 200 | -43*** (Allopurinol) | -47*** (Mannitol) | -50*** (α-tocopherol) |

Furthermore, the effect of these compounds on the nonenzymatic peroxidation of microsomal membrane lipids was also studied. The compounds **6c**, **7b**, and **7d** at 200 µg /ml reduced the microsomal lipid per-oxidation by 35%, 26%, and 23% respectively as compared to α-tocopherol with 50% inhibition

### 4.2 LDL oxidation activity

The cascade of cellular processes can lead to the formation of fatty streaks and ultimately atherosclerotic lesions in the arterial wall as it is triggered by oxidative modification of LDL, so the compounds **6c**, **7b**, and **7d** were also examined for their anti-oxidant activities on human LDL oxidation induced by cupric ions (CuSO_4_). Aerobic oxidation of LDL even in the absence of metal ions caused formation of TBARS (nmol MDA/mg protein), which were greatly increased by 10 to 15 folds in the presence of Cu^+2^. Addition of compounds **6c**, **7b**, and **7d** at 200 µg/ml concentrations in above reaction mixture attenuated LDL oxidation by 32%, 24%, and 17% respectively. (Figure 1)

**Figure 1.**
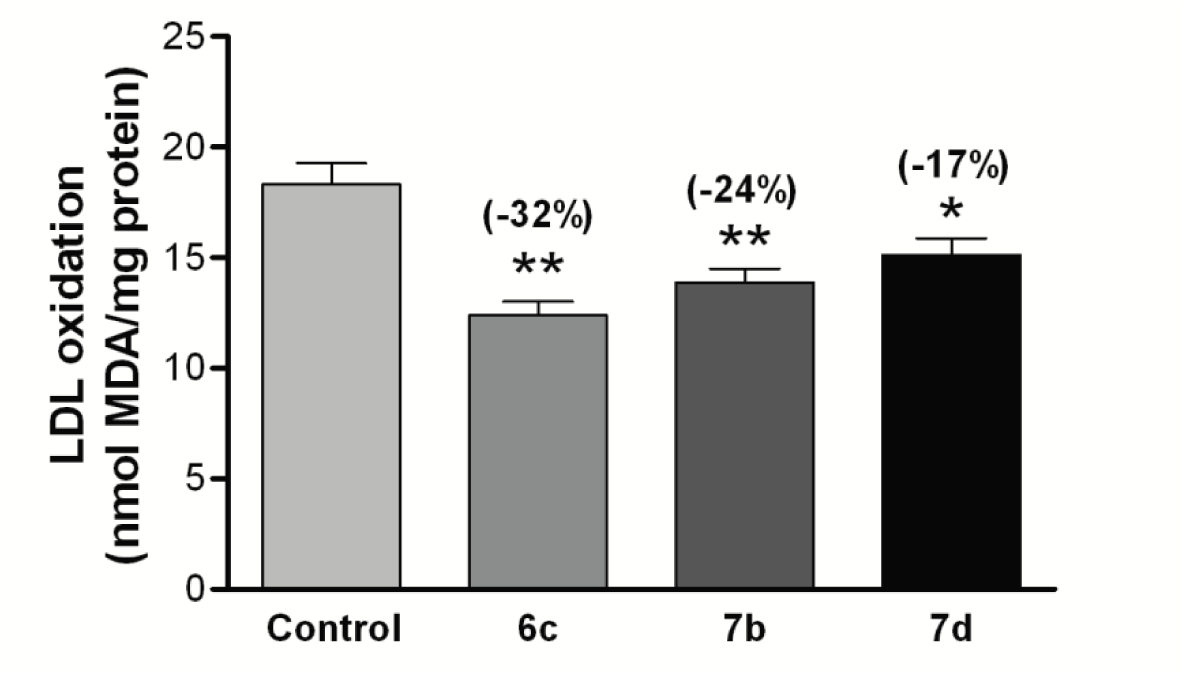
Compounds **7b, 7d** and **6c** reduced the LDL-oxidation at dose of 200µg/ml in normal donor blood sample. Data expressed as mean ± S.D. *P < 0.05; **P < 0.01 compared to treated values with Control.

### 4.3 In vivo anti-dyslipidemic activity

The *in vivo* anti-dyslipidemic activity of these derivatives was also evaluated in triton induced hyperlipidemic model based on the reduction of experimentally induced elevated plasma lipid and MDA levels Table 2 [24–26]. Experimental hyperlipidemia was successfully established 24 h after Triton. WR 1339 administration, with an increase in plasma total cholesterol (3.10 fold), phospholipids PL (3.25 fold), and triglyceride levels (2.67 fold), compared to control. Triton treated rats caused inhibition of plasma PHLA (post heparin lipolytic activity) (38.54 fold) as compared to control.

All the compounds reduced the examined parameters in the plasma of hyperlipidemic rats however, compounds **6c** and **7d** (100 mg/kg b.w) improved the serum lipoproteins TG level in triton induced hyperlipidemic rats by 26% and 24% respectively. The analysis of hyperlipidemic serum of triton administered rats showed significant increase in the level of VLDL-TG and LDL-TG followed by decrease in HDL-TG as compared to control rats.

Treatment with compounds **6c** and **7d** significantly reversed the increased levels of VLDL-TG and LDL-TG and decreased HDL-TG in triton induced hyperlipidemic rats (Figure 2). Under the same experimental conditions, gemfibrozil (used as anti-dyslipidemic drugs), at the same dose,reduced plasma total cholesterol by 33%, PL by 31% and triglycerides by 35 %. Compound **6c** and **7d** also showed inhibition of post heparin. lipolytic activity (PHLA) by 15% and 14%respectively, as compared to gemfibrozil which showed 18% of reversal activity of this enzyme.

**Figure 2.**
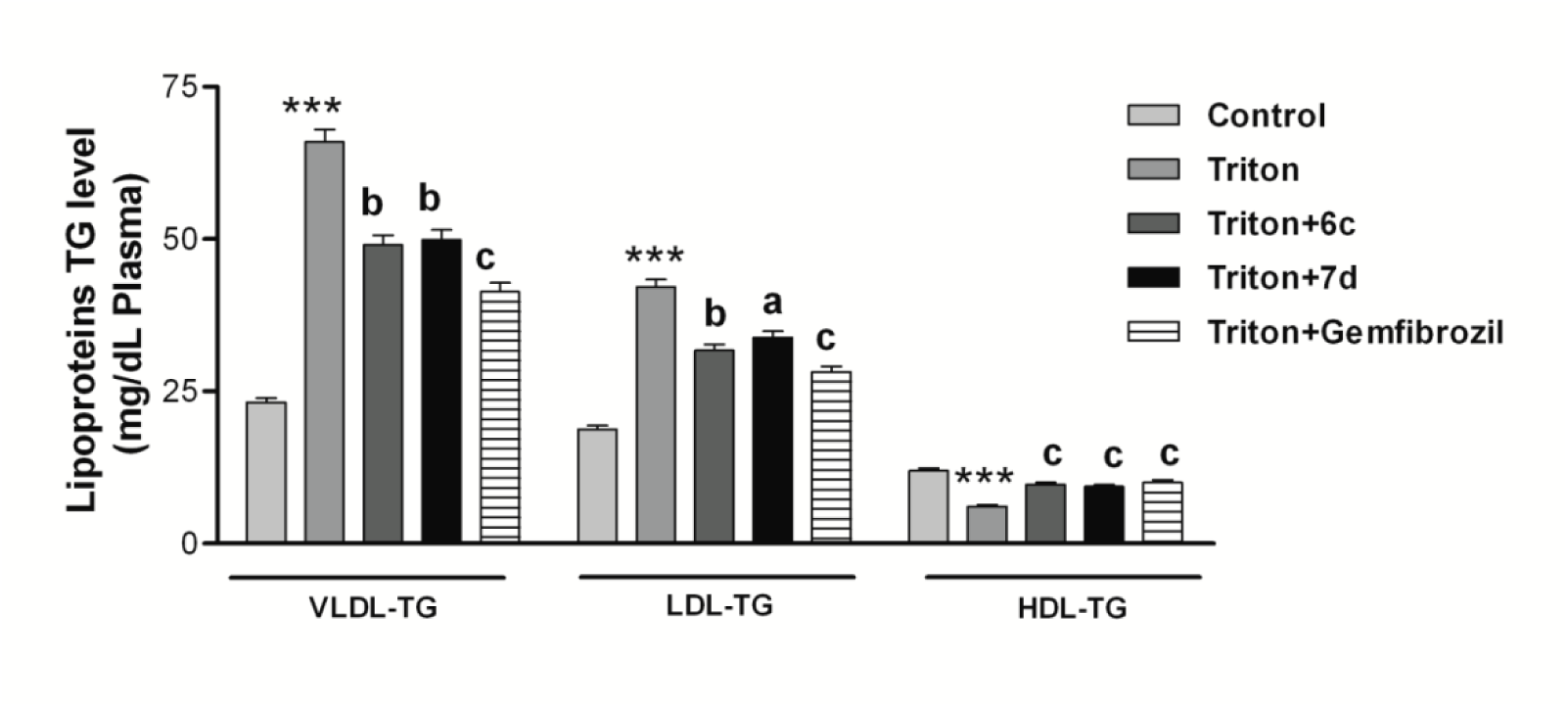
Effect of compound 6c and 7d on lipoprotein triglyceride level in triton induced hyperlipidemic rat. Compound 6c and 7d at dose of 100mg/kg attenuate the level of VLDL-TG and LDLTG followed by increase in HDL-TG level in triton induced hyperlipidemic rats. Each parameter represents pooled data from 6 rats/group and values are expressed as mean ± S.D. ***P < 0.001 between control and triton, ^a^P < 0.05; ^b^P < 0.001; ^c^P < 0.001 between control and triton verses treated rats. Note: Gemfibrozil (100 mg/kg) had taken as standard drug.

| Animal Groups | LIPIDS PROFILE |  |  | PHLA ( nmol of free fatty acids formed/h/ml of plasma) |
| --- | --- | --- | --- | --- |
|  | TC (mg/dl) | PL (mg/dl) | TG (mg/dl) |  |
| **Control** | 78.53±5.13 | 82.64±4.71 | 79.43±6.08 | 18.86±1.37 |
| **Triton** | 244.27±16.84<sup>C</sup><br>(+3.110 Fold) | 269.23±19.82<sup>C</sup><br>(+3.257 Fold) | 239.38±16.38<sup>C</sup><br>(+3.013 Fold) | 11.59±0.93<sup>C</sup><br>(-38.54 Fold) |
| **Triton+5a** | 214.95±17.32* (-12%) | 242.30±17.83* (-10%) | 213.04±17.83* (-11) | 11.93±0.79 NS (+9%) |
| **Triton+5b** | 227.17±12.87 NS (-7%) | 255.76±22.31 NS (-5%) | 220.22±15.39 NS (-8) | 12.51±0.96 NS (+3%) |
| **Triton+5c** | 224.72±18.69 NS (-8%) | 253.07±19.47 NS (-6%) | 220.22±18.46 NS (-8) | 12.63±0.66 NS (+3%) |
| **Triton+5d** | 210.07±14.63* (-14%) | 231.53±18.72* (-14%) | 208.26±16.26* (-13) | 13.67±0.51 NS (+8%) |
| **Triton+6a** | 222.28±29.35 NS (-9%) | 247.69±15.82 NS (-8%) | 217.83±13.29 NS (-9) | 11.93±0.71 NS (+3%) |
| **Triton+6b** | 219.84±17.26* (-10%) | 239.61±13.67* (-11%) | 208.26±18.61* (-13) | 12.05±0.73 NS (+4%) |
| **Triton+6c** | 180.75±13.49*** (-26%) | 196.53±16.11*** (-27%) | 179.53±13.73*** (-25) | 11.93±0.52* (+15%) |
| **Triton+6d** | 217.40±14.83* (-11%) | 242.30±15.48* (-10%) | 210.65±12.96* (-12) | 13.32±1.2 NS (+4%) |
| **Triton+7a** | 212.51±18.27* (-13%) | 242.30±17.91* (-10%) | 210.65±14.52* (-12) | 12.28±0.88 NS (+6%) |
| **Triton+7b** | 202.74±15.91* (-17%) | 226.15±14.26* (-16%) | 203.47±18.21* (-15) | 12.74±0.91* (+10%) |
| **Triton+7c** | 217.40±16.62* (-11%) | 242.30±21.68* (-10%) | 215.44±17.33* (-10) | 12.16±0.63 NS (+5%) |
| **Triton+7d** | 185.64±11.63*** (-24%) | 199.23±16.19*** (-26%) | 184.32±11.85*** (-23) | 13.21±1.11* (+14%) |
| **Triton+Gemfibrozil** | 163.66±11.38*** (-33%) | 185.76±11.66*** (-31%) | 155.59±10.72*** (-35%) | 13.67±0.51* (+18%) |

\*P<0.05; \*\* P<0.01; and \*\*\* P<0.001 is comparison between compounds treated rat vs Triton treated rat

### 4.4 Lipoprotein lipase activity of active compounds

Lipoprotein lipase (LPL) is a major enzyme in overall lipid metabolism and transport, being responsible for hydrolysis of triglycerides present in circulating lipoproteins [27, 28] Decreased adipose tissue LPL activity related to the hypertriglyceridaemia condition which in turn associated with insulin resistance and type II diabetes [29].Therefore, the effect of active compounds **6c** and **7d** were further evaluated for lipoprotein lipase activity (LPLA) in triton induced hyperlipidemic rats. Administration of triton in rats markedly decreases in LPL activity in liver.

**Figure 3.**
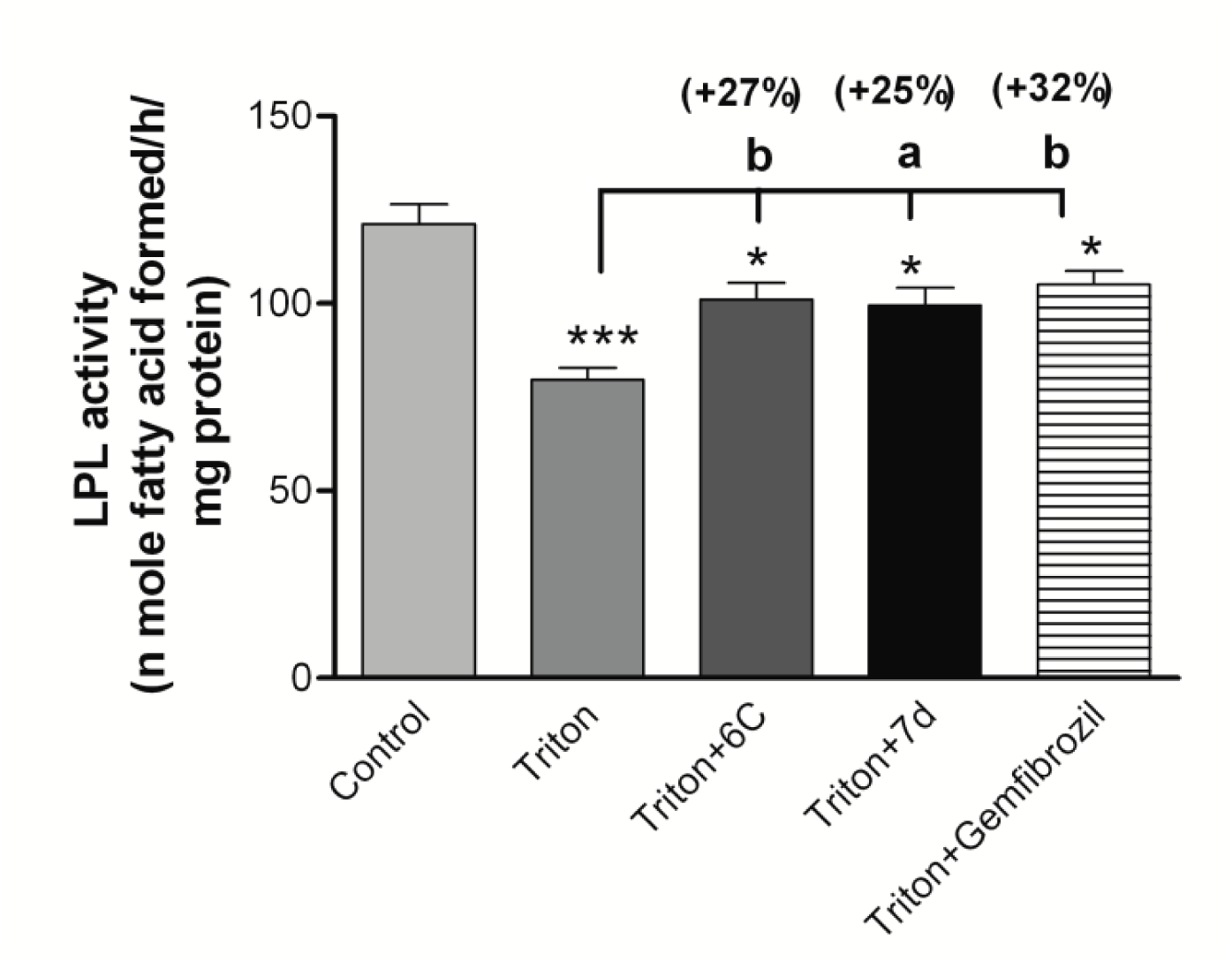
Effect of compound 6c and 7d on lipoprotein lipase activity in triton induced hyperlipidemic rat

### 4.5 Molecular modeling study

The knowledge about the 3D structure of the target may significantly add to the understanding of important interactions which in turn responsible for biological activity variation among the different ligands acting on the same target [29, 30]. The predicted binding mode of compound 6c at LPA binding revealed that the nitrogen atom of indole ring makes H-bond contact with Glu-298 residue. Thr-238 seems to play an important role to the ligand binding through H-bond interaction where it makes H-bond acceptor and donor contact with hydrazine and ketone group of **6c** respectively. These H-bond interactions also provided directionality to the molecule and placed phenyl group at the hydrophobic core made by Pro-241, Phe-239, and Thr-213. Like **6c**, residue Thr-238 played important role in proving anchorage to compound **7d** through H-bond. Indole ring of **7d** occupied hydrophobic pocked through π-cation interaction with Lys-294 whereas on the other end phenyl ring made similar interaction with Arg-214 thus make **7d** energetically favorable at LPA binding site.

### 4.6 Effect of compound **6c** and **7d** on 3T3-L1 cells

Excessive accumulation of adipose mass tissue not only contributes to obesity but also increase the risk of cardiovascular disease and type-2 diabetes. Therefore, the active compounds of the series **6c** and **7d** were also evaluated on 3T3-L1 preadipocytes. To investigate the anti-adipogenic effect of compound **6c** and **7d**, cells were treated with and without MDI adipogenic media and 20µM of compound **6c** and **7d** dissolved in MDI adipogenic media (MDI is adipogenic media with IBMX, dexamethosone and Insulin used to differentiate preadipocytes into adipocytes). Adipogenesis was assessed by oil red O (ORO) staining of lipid droplets. ORO staining results showed that Compound **6c** and **7d** reduced MDI mediated lipid accumulation at 20µM as observed microscopically (Figure 4A). MDI-induced greater lipid accumulation was reduced approximately 44% and 31% in the cells treated with compound **6c** and **7d** at 20µM, as measured spectrophotmetrically (Figure 4B).

**Figure 4.**
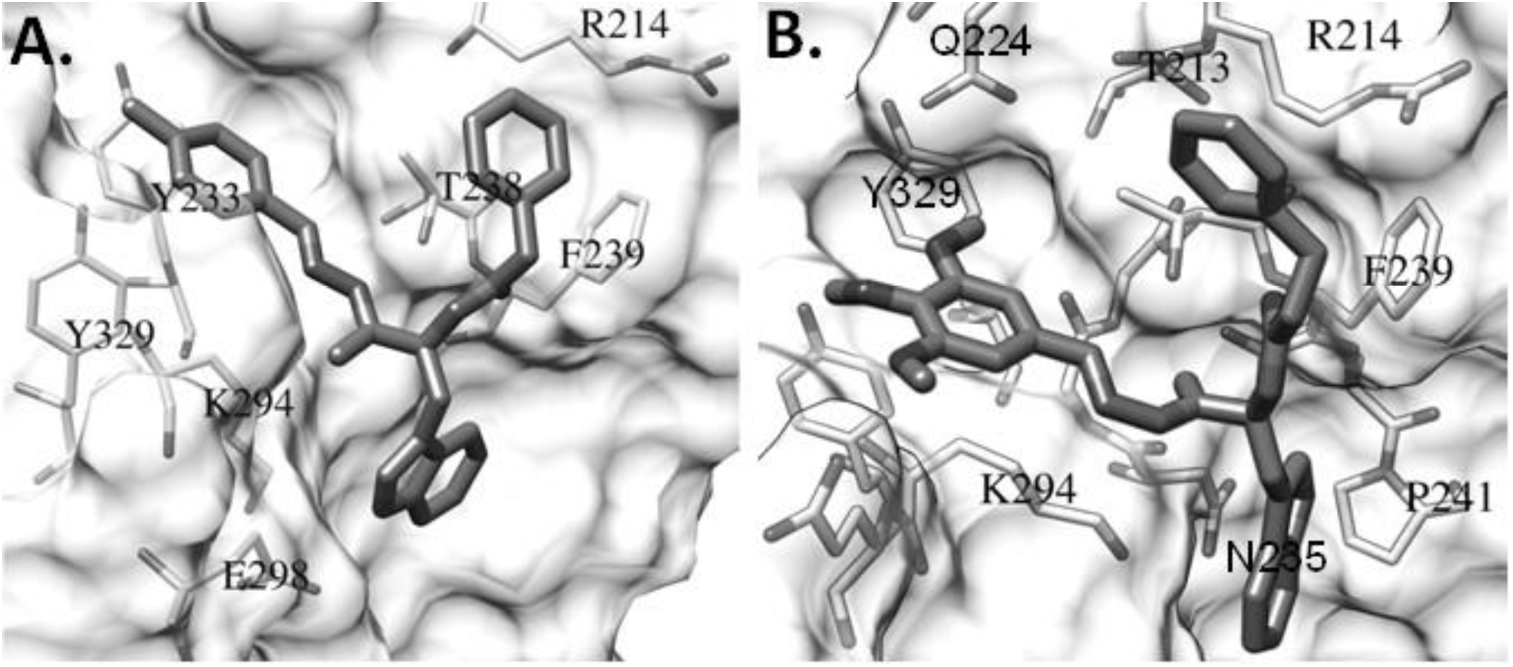
Lowest energy binding poses of compound 6c (A) and 7d (B) docked onto human LPA protein.

**Figure 5.**
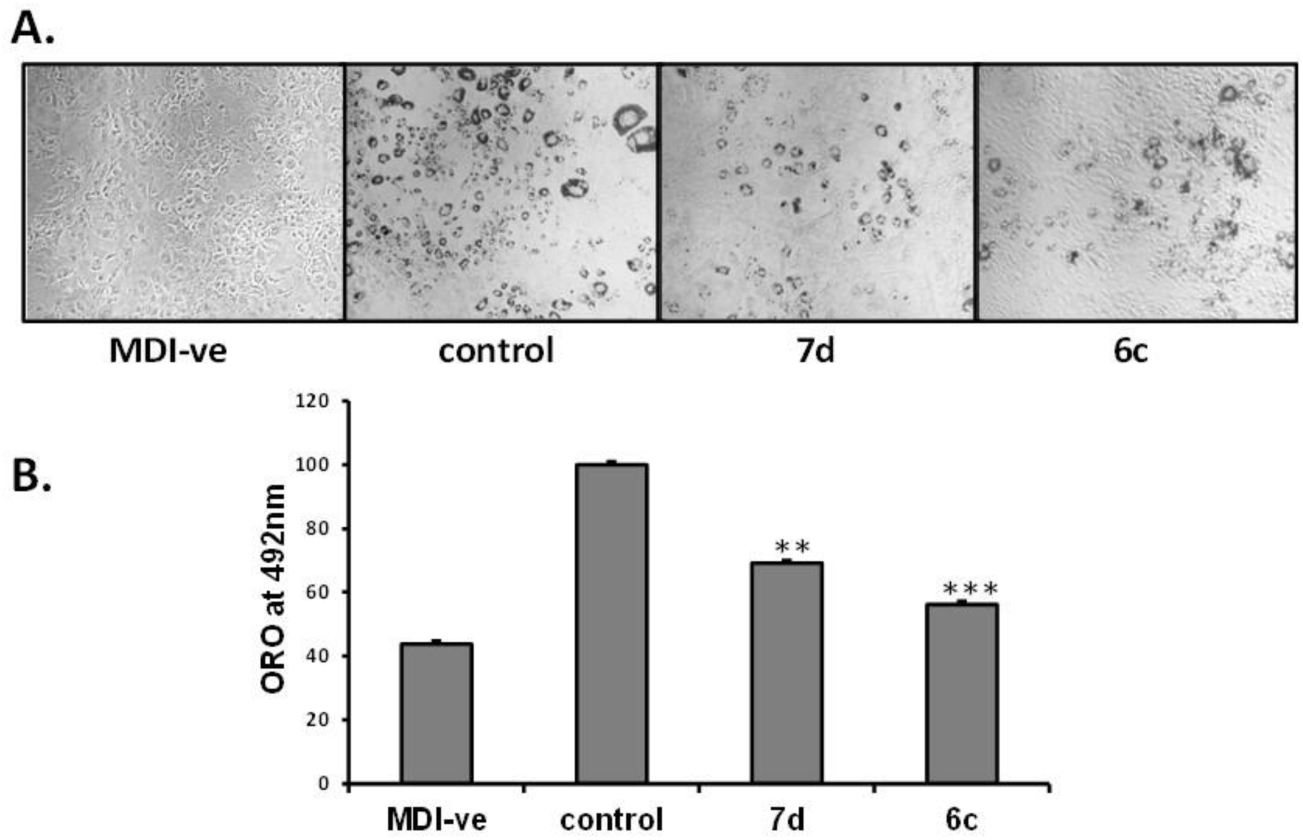
(A and B). Effects of compound **6c** and **7d** on MDI media induced adipogenesis in 3T3-L1 preadipocytes, compound **6c** and **7d** inhibited differentiation of 3T3-L1 preadipocytes as indicated by staining with Oil Red O solution. Data are the mean ± S.D., n=3 and *p<0.001* when compared with MDI induced control group.

### 4.7 In vivo Anti-hyperglycemic activity

Taking into consideration the promising anti-oxidant, anti-dyslipidemic and anti-adepogenic activity of compound **6c** and **7d**, these compounds were further tested for their anti-diabetic activity. Figure 5a and 5b shows the effect of compounds **6c**, **7d** and standard drug on decline in blood glucose level on streptozotocin-induced diabetic rats at various time intervals. Metformin was taken as positive control. It is evident from the results that both compounds showed singnificant decline in blood glucose levels on streptozotocin-induced diabetic rats. However, compound **6c** was found to be the most active compound of the series. Compounds **6c**, **7d** and Standard drug metformin demonstrated maximum decline in blood glucose to the tune of 20.5 % (p<0.01) 16.7 % (P<0.05) and 23.3 % (p<0.01) at 5 h and 24.3 % (p<0.01), 18.2 % (p<0.01)and 29.5 % (p<0.01) at 24 h respectively on the STZ-induced diabetic rats at 100 mg/kg oral dose.

**Figure 5.**
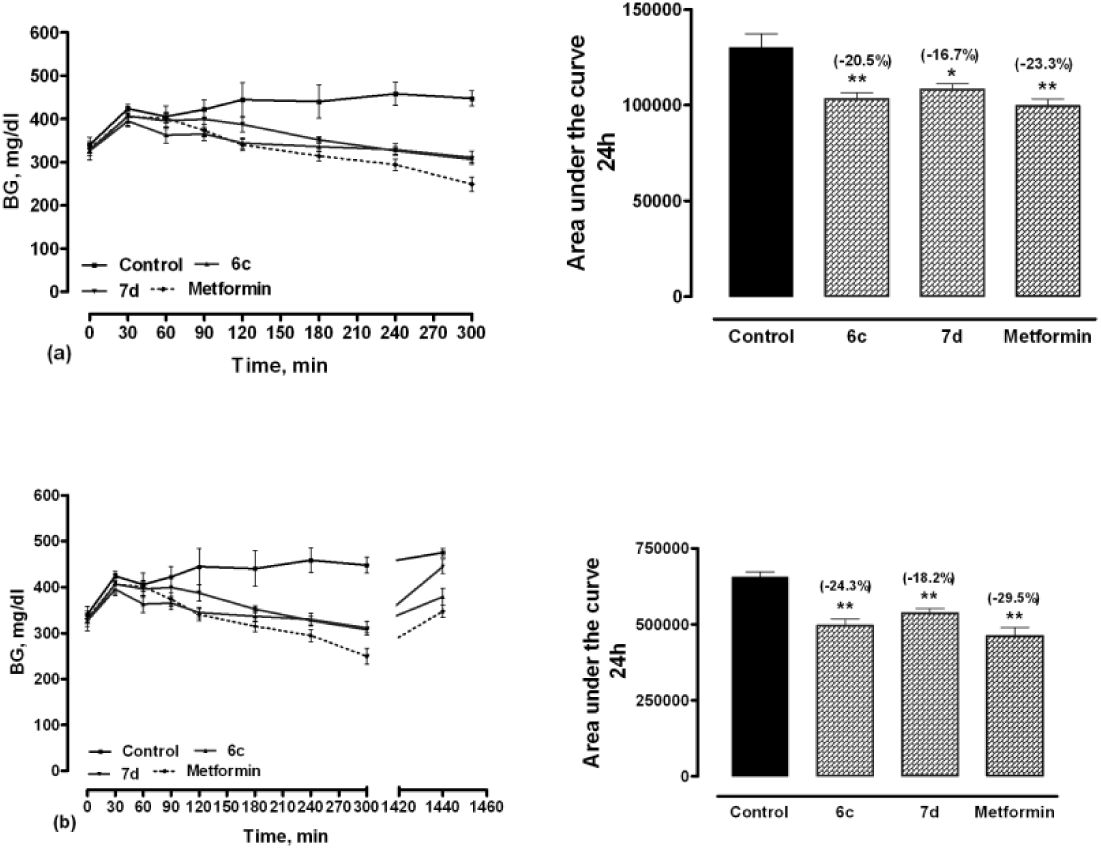
Effect of test compounds on blood glucose profile of streptozotocin-treated diabetic rats. Statistical analysis was made by Dunnet test (Prism Software 3)

## Conclusion

In summary, a series of 12 substituted hydrazone derivatives were synthesized and evaluated for the anti-oxidant, anti-dyslipidemic activity in an acute triton induced hyperlipidemic model. Among the 12 compounds two compounds **6c** and **7d** showed good anti-oxidant activity with enhanced protection against lipid peroxidation. Both the compounds **6c** and **7d** also showed effective protection against LDL oxidation and reduced the TC and TG levels by 23-26% with improved lipoprotein lipase activity corroborated by the binding energies of these compounds at lipoprotein lipase binding site. Molecular docking study also suggested important role of Thr-238 to the ligand binding at lipoprotein lipase active site. Both the compounds **6c** and **7d** also showed anti-adipogenic and anti-hyperglycemic activity, where compound **6c** was found to be the most active compound with 44% reduction in lipid accumulation and also reduce the blood glucose by 20.5% at 5h and 24.3% at 24h in comparison to standard drug metformin. Thus, these compounds with balanced activities may be useful in the treatment of metabolic disorder.

## 6 Characterization details

### 6.1 Chemistry

Melting points were determined on an electrical heated m. p. apparatus /using silicon oil bath. Reactions were monitored by thin layer chromatography on self-made plates of silica gel G (Merck, India) or 0.25mm ready-made plates of silica gel 60F254, (Merck, Darmstadt, Germany). Column chromatography was performed on silica gel (Merck, 60 to 120mesh). Infrared spectra (IR) were recorded on Perkin-FTIR model PC spectrophotometer with frequency of absorptions reported in wave numbers. MS were recorded on JEOL spectrometer with fragmentation pattern reported as values. 1H NMR was recorded on Bruker spectrometer with a multinuclear inverse probe head with gradient at room temperature (298 K) using CDCl3 as solvent and tetramethylsilane (TMS) as internal standard. Chemical shifts were described in parts per million (ppm) relative to TMS (0.00 ppm) using scale, and coupling constants were reported in hertz (Hz).

#### 6.1.1 Synthesis of DL-tryptophan methyl ester hydrochloride

Thionyl chloride (7ml) was added drop wise to a stirred suspension of DL-tryptophan 1 (10g) in dry methanol (45 ml) at −10°C during 30 min and stirring continued for 6 hrs during which temperature was gradually raised from −10°C to room temperature (25°C). The reaction mixture was filtered under suction and afforded the cream colored product tryptophan methyl ester which was washed with ether (10ml.) and dried.Yield 9.7 gm (92%); m.p: 220-223°C; [α]^D^_20_ 0; ^1^HNMR (200 MHz, CDCl3+ DMSO-d6): δ 3.45 (d, J=5.90, 2H), 3.75 (s, 3H), 4.17 (m, 1H), 7.07(m, 2H), 7.28(s, 1H), 7.41(d, J=7.72, 1H), 7.52(d, J=7.54, 1H), 8.60 (brs, 1H), 10.76(s,1H); IR (KBr): 603, 731, 1034,1225, 1285, 1352, 1443, 1499, 1580, 1748, 2021, 3292; MS(ESI+): m/z 218 (M+).

#### 6.1.2 Synthesis of DL-N-benzoyl tryptophan methylester (2)

Benzoyl chloride (1.15ml) in dry THF (5ml) was added to a stirred solution of dl tryptophan methyl ester (**1)** (2.18gm, 1eq) and dry triethylamine (1.66 ml, 1.2eq) in dry THF (8ml) during 15 min and was allowed to stir for 2 hrs at room temperature. The reaction mixture was concentrated under vacuum; the residue was triturated with water (20 ml) to get 2b, which was crystallized with methanol. Yield 2.96 gm (92%); m.p: 111°C; [α]^D^_20_0; ^1^HNMR (300 MHz, CDCl3+ DMSO-d6): δ 3.21 (m,1H), 3.46 (m, 1H), 3.68(s,3H), 4.81 (m, 1H), 7.11-7.18 (m, 3H), 7.28(s, 1H), 7.61-7.67(m, 3H), 7.97(m, 2H), 9.57 (s, 1H), 10.85 (s,1H); MS(ESI+): m/z 323 (M+1)+. Compounds 3–4 were synthesized in the same manner.

#### 6.1.3 Synthesis of (DL)-N-(1-hydrazinyl-3-(1H-indol-3-yl)-1-oxopropan-2-yl) benzamide (5)

A solution of 2b (1.7 g, 5.5 mmol) in ethanol (20 mL) containing 85% hydrazine hydrate (10 mL) was refluxed for 6 h. The reaction mixture was concentrated under vacuum, to give white solid which was triturated with water and dried to give compound 5, which was further crystallized with methanol. Yield 2.2 gm (70%); m.p: 225-228°C; [α]^D^_20_0^1^HNMR (300 MHz, CDCl3+ DMSO-d6): δ 2.91-2.96 (m,1H), 3.09-3.16 (m, 1H), 3.92 (s, 2H), 4.03-4.08 ( m, 1H), 6.87-7.34 (m, 8H), 7.85 (m, 2H), 8.34(s, 1H), 9.06(s, 1H), 10.23(s, 1H); MS(ESI+): m/z 323 (M+1).

Compounds 6–7were synthesized in the same manner.

#### 6.1.4 Synthesis of N-(1-(2-( 4-chlorobenzylidene) hydrazinyl)-3-(1H-indol-3-yl)-1-oxopropan-2-yl)benzamide (5a)

Equimolar quantities of hydrazide (**5**) and the appropriate aldehyde existing in the liquid or solid state under normal conditions, were mixed in DMSO. The mixtures were stirred at room temperature for 4 hrs to give white solid compound (**5a**). Yield: 0.154 g (70.0 %); m.p.: 132°C; 1H NMR (300 MHz, CDCl3): δ 3.01-3.16 (m, 1H), 4.72-4.78 (m, 1H), 5.58-5.61(m,1H), 6.70-7.67 (m, 14H), 7.82 (s, 1H), 8.53 (s, 1H), 9.75 (s, 1H), 11.01 (s, 1H); FTIR (KBr): cm-1 754,1215, 1484, 1532, 1664, 2998, 3341,; ESI-Ms: m/z (M+1)+ 445; Anal. Calcd for C25H21ClN4O2: C, 67.49; H, 4.76; N, 12.59; found: C, 67.18; H, 4.41; N, 12.34 %

#### 6.1.5 N-(3-(1H-indol-3-yl)-1-(2-(4-methoxybenzylidene) hydrazinyl)-1-oxopropan-2-yl) benzamide (5b)

Yield: 0.157 g (72.0 %); m.p.: 200°C; 1H NMR (300 MHz, CDCl3): δ 3.40-3.45 (m,1H), 3.89 (s, 3H), 5.01-5.03 (m, 1H), 5.86-5.89 (m,1H), 6.88-7.91 (m, 14H), 7.98 (s, 1H), 8.60 (s, 1H), 9.78 (s, 1H), 10.91 (s, 1H); FTIR (KBr): cm-1 759, 1216, 1456, 1517, 1625,1661, 2928,3385; ESI-Ms: m/z (M+1)+ 441; Anal. Calcd for C26H24N4O3: C, 70.89; H, 5.49; N, 12.72; Found: C, 70.68; H, 5.41; N, 12.84 %.

#### 6.1.6 N-(3-(1H-indol-3-yl)-1-(2-(4-methylbenzylidene) hydrazinyl) -1-oxopropan-2-yl)benzamide (5c)

Yield: 0.154 g (69.0%); m.p.: 190°C; 1H NMR (300 MHz, CDCl3): δ 2.84(s,3H), 3.36-3.40 (m,1H), 4.98-5.03 (m, 1H), 5.82-5.86(m,1H), 70.8-8.11 (m, 14H), 7.90 (s, 1H), 8.56 (s, 1H), 10.18 (s, 1H), 11.01 (s, 1H); FTIR (KBr): cm-1 767, 1218, 1456, 1517, 1632,1665, 2928,3375; ESI-Ms: m/z (M+1)+ 425; Anal. Calcd for C_26_H_24_N_4_O_2_: C, 73.56; H, 5.70; N, 13.20; Found: C, 73.48; H, 5.46; N, 13.04 %.

#### 6.1.7 (E)-N-(3-(1H-indol-3-yl)-1-oxo-1-(2-(3,4,5-trimethoxybenzylidene) hydrazinyl)propan-2-yl)benzamide (5d)

Yield: 0.207 g (74.9 %); m.p.: 120 °C; ^1^H NMR (300 MHz, CDCl_3_): *δ* 3.43-3.48 (m,1H), 3.89 (s, 9H), 5.04-5.07 (m, 1H), 5.93-5.96 (m,1H), 6.94-7.89 (m, 12H), 7.94 (s, 1H), 8.59 (s, 1H), 9.59 (s, 1H), 10.94 (s, 1H); FTIR (KBr): cm^-1^ 760, 1217, 1460, 1517, 1632,1676, 3397; ESI-Ms: ^m/z (M+1)^+^ 501; Anal. Calcd for C_28_H_28_N_4_O_5_: C, 67.19; H, 5.64; N, 11.10; found: C, 66.48; H,^ 5.56; N, 11.04 %.

#### 6.1.8 N-(1-(2-(4-chlorobenzylidene)hydrazinyl)-3-(1H-indol-3-yl)-1-oxopropan-2-yl)-2-phenylacetamide (6a)

Yield: 0.196 g (72.0 %); m.p.: 170 °C; 1H NMR (300 MHz, CDCl3): δ 2.58 (s,2H), 3.51-3.57 (m,1H), 4.77-4.79 (m, 1H), 5.64-5.66 (m, 1H) 6.89-7.89 (m, 14H), 8.02 (s, 1H), 8.61 (s, 1H), 9.93 (s, 1H), 11.13 (s, 1H); FTIR (KBr): cm-1 758, 1217, 1486, 1517,1651, 3401; ESI-Ms: m/z (M+1)+ 459; Anal. Calcd for C_28_H_28_N_4_O_5_: C, 68.04; H, 5.05; N, 7.72; found: C, 67.88; H, 4.96; N, 7.44 %.

#### 6.1.9 N-(3-(1H-indol-3-yl)-1-(2-(4-methoxybenzylidene)hydrazinyl)-1-oxopropan-2-yl)-2-phenylacetamide (6b)

Yield: 0.234 g (69.0 %); m.p.: 175 °C; 1H NMR (300 MHz, CDCl3): δ 2.60 (s,2H), 3.01-3.29 (m, 1H), 3.82 (s, 3H), 4.80-4.83 (m, 1H), 5.65-5.70 (m,1H), 6.64-7.84 (m, 14H), 7.92 (s, 1H), 8.58 (s, 1H), 9.85 (s, 1H), 10.91 (s, 1H); FTIR (KBr): cm-1 760, 1217, 1420, 1510,1668, 3399; ESI-Ms: m/z (M+1)+ 455; Anal. Calcd for C_27_H_26_N_4_O_3_: C, 71.35; H, 5.77; N, 12.33; found: C, 70.88; H, 5.66; N, 12.13 %.

#### 6.1.10 N-(3-(1H-indol-3-yl)-1-(2-(4-methylbenzylidene)hydrazinyl)-1-oxopropan-2-yl)-2-phenylacetamide (6c)

Yield: 0.290 g (75.0 %); m.p.: 180°C; 1H NMR (300 MHz, CDCl3): δ 2.64 (s, 2H), 3.13( s,3H), 3.54-3.57 (m,1H), 4.81-4.83 (m, 1H), 5.69-5.70 (m,1H), 6.91-7.52 (m, 12H), 7.55-7.89 (m,2H), 7.99 (s, 1H), 8.59 (s, 1H), 10.13 (s, 1H), 11.13 (s, 1H); FTIR (KBr): cm-1 760, 1215, 1421, 1521,1643, 3406; ESI-Ms: m/z (M+1)+ 439; Anal. Calcd for C27H26N4O2: C, 73.95; H, 5.98; N, 12.78; Found: C, 73.83; H, 5.72; N, 12.53 %.

#### 6.1.11 N-(3-(1H-indol-3-yl)-1-oxo-1-(2-(3,4,5-trimethoxybenzylidene)hydrazinyl)propan-2-yl)-2-phenylacetamide (6d)

Yield: 0.318 g (69.0 %); m.p.: 160 °C; ^1^H NMR (300 MHz, CDCl_3_): *δ* 2.58 (s, 2H), 3.54-3.57 (m,1H), 3.86(s,9H), 4.91-4.93 (m, 1H), 5.67-5.69 (m,1H), 7.11-7.52 (m, 12H), 8.19 (s, 1H), 8.69 (s, 1H), 10.11 (s, 1H), 11.03 (s, 1H); FTIR (KBr): cm^-1^ 759, 1221, 1414, 1502,1652, 3406; ESI-Ms: m/z (M+1)^+^ 515; Anal. Calcd for C_29_H_30_N_4_O_5_: C, 67.69; H, 5.88; N, 10.89; found: C, 67.56; H, 5.85; N, 10.23 %.

#### 6.1.12 N-(1-(2-(4-chlorobenzylidene)hydrazinyl)-3-(1H-indol-3-yl)-1-oxopropan-2-yl)-3-phenylpropanamide (7a)

Yield: 0.191g (71.0 %); m.p.: 205 °C; 1H NMR (300 MHz, CDCl3): δ 2.47-2.49 (m, 2H), 2.83-2.86( m,2H), 3.15-3.17 (m,1H), 4.81-4.83 (m,1H), 5.54-5.63 (m, 1H), 6.91-7.64 (m, 14H), 7.93 (s, 1H), 8.13 (s, 1H), 10.48 (s, 1H), 11.28 (s, 1H); FTIR (KBr): cm-1 762, 1216, 1421, 1510,1653, 3403; ESI-Ms: m/z (M+1)+ 473; Anal. Calcd for C_27_H_25_ClN_4_O_2_: C, 68.56; H, 5.33; N, 11.85; found: C, 68.16; H, 5.05; N, 11.53 %.

#### 6.1.13 N-(3-(1H-indol-3-yl)-1-(2-(4-methoxybenzylidene)hydrazinyl)-1-oxopropan-2-yl)-3-phenylpropanamide (7b)

Yield: 0.22 g (72.6 %); m.p.: 197 °C; 1H NMR (300 MHz, CDCl3): δ 2.44-2.52 (m, 2H), 2.80-2.85( m,2H), 3.11-3.14 (m,1H), 3.83 (s,3H), 4.80-4.82 (m,1H), 5.64-5.72 (m, 1H), 7.11-7.84 (m, 14H), 7.95 (s, 1H), 8.15 (s, 1H), 10.50 (s, 1H), 11.18 (s, 1H); FTIR (KBr): cm-1 759, 1216, 1423, 1508,1653, 3341, 3403; ESI-Ms: m/z (M+1)+ 469; Anal. Calcd for C_28_H_28_N_4_O_3_: C, 71.78; H, 6.02; N, 11.96; found: C, 71.56; H, 6.05; N, 11.87 %.

#### 6.1.14 N-(3-(1H-indol-3-yl)-1-(2-(4-methylbenzylidene)hydrazinyl)-1-oxopropan-2-yl)-3-phenylpropanamide (7c)

Yield: 0.264 g (68.3 %); m.p.: 198°C; 1H NMR (300 MHz, CDCl3): δ 2.50-2.53 (m, 2H), 2.63 (s, 3H), 2.85-2.90 ( m,2H), 3.18-3.21 (m,1H), 4.77-4.79 (m,1H), 5.69-5.71 (m, 1H), 6.91-7.79 (m, 14H), 7.90 (s, 1H), 8.05 (s, 1H), 10.22 (s, 1H), 11.19 (s, 1H); FTIR (KBr): cm-1 759, 1215, 1424, 1508,1637, 3380; ESI-Ms: m/z (M+1)+ 453; Anal. Calcd for C_28_H_28_N_4_O_2_: C, 74.31; H, 6.24; N, 12.38; found: C, 74.16; H, 6.11; N, 12.17 %.

### 6.1.15 (N-(3-(1H-indol-3-yl)-1-oxo-1-(2-(3,4,5-trimethoxybenzylidene)hydrazinyl)propan-2-yl)-3-phenylpropanamide (7d)

Yield: 0.304 g (71.0 %); m.p.: 190 °C; 1H NMR (300 MHz, CDCl3): δ 2.45-2.50 (m, 2H), 2.83-2.86( m,2H), 3.15-3.18(m,1H), 3.83 (s, 9H), 4.71-4.76 (m,1H), 5.63-5.65 (m,1H), 6.95-7.87 (m, 12H), 7.95 (s, 1H), 8.07 (s, 1H), 10.36 (s, 1H), 11.15 (s, 1H); FTIR (KBr): cm-1 757, 1217, 1418, 1504,1664, 3396; ESI-Ms: m/z (M+1)+ 529; Anal. Calcd for C_30_H_32_N_4_O_5_: C, 68.17; H, 6.10; N, 10.60; found: C, 67.99; H, 6.01; N, 10.57 %.

## Acknowledgements

We thank Sophisticated Analytical Instruments Facility, CDRI for providing analytical data. The authors are also thankful for financial assistance by Council of Scientific and Industrial Research (CSIR) (KV, RS, SV, AM). AKG is supported by a fellowship from the Gulf Coast Consortia, on the Computational Cancer Biology Training Program (CPRIT Grant No. RP170593). The CDRI communication number allotted to this manuscript is XXXX.

